# Could cyanobacteria have made the salinity transition during the late Archean?

**DOI:** 10.1101/216416

**Authors:** A. Herrmann, M.M. Gehringer

**Affiliations:** Department of Plant Ecology and Systematics, Faculty of Biology, Technical University of Kaiserslautern, 67663 Kaiserslautern, Germany

**Author notes:** &.

**Keywords:** Archaean, Cyanobacteria, salinity transition, net photosynthesis, pigments, *Chroococidiopsis thermalis*, *Gloeobacter violaceus*

## Abstract

Modern molecular evolutionary studies suggest the freshwater origin of cyanobacteria during the late Archean, about 2.7 Ga ago, with an even earlier evolution of oxidative photosynthesis. The large amount of oxygen required to oxygenate the Earth’s atmosphere during the Great Oxygenation Event (GOE) ~2.4 Ga is thought to have been produced by large cyanobacterial blooms in the open ocean. This then poses the question as to whether ancient lineages of cyanobacteria would have survived the salinity transition. This study investigates the effect of increasing salinity on the photosynthetic efficiency of two modern day descendants of ancient cyanobacteria, *Chroococidiopsis thermalis* PCC7203 and the root species, *Gloeobacter violaceus* PCC7421. Organisms were cultured in fresh, brackish or sea water analogous media under a present atmospheric level (PAL) atmosphere or an atmosphere with reduced O_2_ and elevated CO_2_ (rO_2_eCO_2_). The net photosynthesis (NP) rates were determined in liquid cultures, while the O_2_ and redox profiles were determined in pseudomats. While *C. thermalis* PCC7203 was able to grow under increasing salinities under both atmospheres tested, *G. violaceus* PCC7421 could not make the salinity change to sea water. NP rates were reduced for *C. thermalis* under increasing salinities, as were the levels of dissolved O_2_ in the media. A gene screen indicated that *C. thermalis* genome carries genes for both sucrose and trehalose synthesis, whereas *G. violaceus* has only the later genetic component, suggesting a mechanism for their differing salt tolerances. This study supports the hypothesis of Cyanobacterial evolution in freshwater environments and their transition into increasingly salty environments during the late Archaean, prior to the GOE.

## Introduction

The early origin of life is often assumed to have evolved in the deep oceans as this is where the first geochemical records and fossils were identified. Modern evolutionary and geochemical studies, however, indicate that widespread shallow water marine (Heubeck *et al*., 2016, Homann *et al*., 2016) and non-marine life was well established in the late Archean (Stueken *et al*., 2015, Djokic *et al*., 2017). Furthermore, strong sedimentary evidence exists for terrestrial land biota (Lenton & Daines, 2017). While the first mats were presumably powered by anoxic phototrophs, oxidative phototrophs would have exhibited higher primary productivity and therefor capable of leaving the clear traces we observe in the geological records today. Cyanobacteria are the pioneer species of modern day biological soil crusts (Büdel *et al*., 2014) and have demonstrated their diverse adaptability to inhabit environments from hot springs to dry Antarctic mats. The earliest biological mats, whether in fresh or marine shallow water environments, or subaerial, would have been dependent on the primary productivity of early cyanobacteria to provide both biological accessible C and N.

Recent studies based on the analysis of sequenced cyanobacterial genomes places the origin of cyanobacteria at ~2.7 Ga in a freshwater environment (Sánchez-Baracaldo *et al*., 2014, Sanchez-Baracaldo *et al*., 2017). These studies identified modern day descendents of cyanobacteria that are the most closely related to the cyanobacteria proposed to have lived during the Archean. At the root of the tree is *Gloeobacter violaceus* PCC7421 and *Gloeobacter kilauensis*, both unicellular freshwater species. *G. kilauensis* is a very slow growing organism and unsuited to largescale laboratory investigations (Saw *et al*., 2013). These strains are of particular interest as they do not contain thylakoids and therefor carry the photosynthetic apparatus on the cell membrane. As the result of this presumed ancient structure, *G. violaceus* keeps tight control of its light harvesting pigments, producing low levels of chlorophyll *a* and phycobilliproteins (Guglielmi *et al*., 1981). *Chroococcidiopsis thermalis* PCC7203, while not located in the cyanobacterial basal clade, is still an anciently derived cyanobacterial species often proposed as an example of the earliest cyanobacteria (Fewer *et al*., 2002) as they are cryptoendolithic (Budel *et al*., 2004) and able to withstand desiccation and raised salinities (Bahl *et al*., 2011, Cumbers & Rothschild, 2014). It has high UV tolerance, is able to withstand harsh desiccation conditions by virtue of its production of a thick exopolymeric substance (EPS) and thick cell walls. Based on these studies, *C. thermalis* PCC7203 has been proposed as an early Earth terrestrial coloniser and also used as a space coloniser (Billi *et al*., 2011, Cockell *et al*., 2011).

It is against this background that we undertook a study to assess the nett photosynthesis activities of modern day descendants of ancient Cyanobacteria, *Chroococidiopsis thermalis* PCC7203 and *Gloeobacter violaceus* PCC7421, to grow under increasing salinities under a modern day atmosphere or an atmosphere representative of the Archean.

## Materials and Methods

### Cultures

Cultures of *Gloeobacter violaceus* PCC7421 and *Chroococcidiopsis thermalis* PCC 7203 were purchased from the Pasteur Culture Collection and maintained in BG11 (PCC) media on a day night cycle of 16:8 hrs in a Percival culture chamber (E-22L). Daylight illumination was supplemented with red light and far red light LED banks at 20 and 60 µmols photons m^−2^ s^−1^ respectively.

#### Growth curve

*C. thermalis* PCC7203 and *G. violaceus* PCC7421 were used in growth curve experiments to compare the effects of increased salinities and altered atmospheres on their growth. Early stationary phase cultures were used to inoculate media (BG11, BG11:ASNIII & ASNIII as per PCC) at an OD_600_ of 0.1 and pipetted in 200 µl volumes into 5 wells of a 96 well tissue culture plate and incubated at 20 µmol photons m^2^ s^−1^ for *G. violaceus* and 60 µmol photons m^2^ s^−1^ for *C. thermalis*. A duplicate plate was incubated in an airtight culture chamber that was gassed daily with rO_2_eCO_2_ gas at 600 and 1000 ppm respectively. Cultures were briefly (~10 min.) exposed to PAL conditions every second day when their optical density at 650 nm (OD_650_) was measured in a microtiter plate reader (Multiscan FC, ThermoFisher Scientific, USA) after 30 s of shaking.

#### Nett photosynthesis experiment

Cultures were grown at 24 °C at 65% humidity in ventilated T_75_ suspension culture flasks (Sarsted, Germany). Large scale cultures (T_175_), in early stationary phase, were split into 2 sets of triplicates, drained of medium and resuspended in media representing one of the following conditions: fresh (BG11), brackish (BG11:ASNIII, 1:1) and salty (ASNIII) culture conditions, with salinities of 1, 16 and 29,5 psu (pratical salinity units) respectively (conductivity measurements; inoLab Cond720, WTW, Germany). The 50 ml cultures in T_75_ ventilated flasks were placed in airtight culture containers with holes (PAL ~410 ppm CO_2_) or rubber stoppers (rO_2_eCO_2_), the latter gassed daily a premixed gas of 600 ppm O_2_ and 1000 ppm CO_2_ with the rest composed of N_2_ gas, at a flow rate of 0.5 L min^−1^ for 15 minutes, and incubated as above.

Samples for photosynthesis were harvested on day 14 (late exponential/early stationary phase) in their appropriate atmospheres, and filtered onto a mixed cellulose filter of 3.0 µm (Millipore SWPP02500, Germany). Filters were maintained on fresh moist agar plates at the appropriate atmosphere until NP assessment. The filters were placed in a sample lichen cuvette of the GFS3000 (WALZ, Germany) and the NP was determined over a range of light intensities. After the material was used for NP determinations, the filters were placed in a 2 ml brown plastic vial to which 100 mg 0.1 mm Silica beads and 1.5 ml 90% neutralized methanol were added. Samples were bead beaten (Retsch) at 35 beats per second for 45 seconds and stored overnight at 4°C. The following morning samples were vortexed and centrifuged for 15 min at 10 000 g. The absorbance of the cleared supernatant was as measured at 665 nm against a blank of 90% neutralized method and the chlorophyll *a* content determined (Meeks & Castenholz, 1972). NP was expressed per µg Chl *a*.

Media was assessed for dissolved Ci using the program CO2SYS, version 2.3 (Lewis et al., 1998) with the experimentally determined values for pH and total alkalinity (TA) as input data. TA was measured using the open-cell titration method described in Dickson *et al*. (2007). The titration was conducted manually with a 10 ml burette (Schott AG, Germany) and recorded with the pH meter model PHM210 (HyXo Oy, Finnland) with a SJ223 pH electrode (VWR, USA) which was also used to determine the input pH after incubation. For the TA calculation the program RStudio version 1.0.153 with the package seacarb was used. The phosphorus content of the samples was assessed photometrically as phosphomolybdenum blue (McKelvie *et al.,* 1995)

### Pigment determinations

Prior to NP determinations, 4 ml of culture was pelleted in each of 2 brown 2 ml centrifuge tubes (repeated centrifugation) and used to determine the chlorophyll a or phycobiliprotein content per ml of culture. Chlorophyll determinations proceeded as above. Phycobilliproteins were extracted using 1.5 ml of 0.1 M potassium phosphate buffer pH 6.8, bead beating and 2 cycles of rapid freeze thawing in liquid nitrogen. Both pigment extractions were left overnight at 4°C before vortexing and centrifugation (15 min at 10 000 g). Chlorophyll a extractions were measured as above. The total phycobilliprotein content was determined from the absorption readings at 562, 615 and 652 nm (Bennett & Bogorad, 1971, Bennett & Bogorad, 1973).

#### Pseudomat experiment

Late stationary phase cultures were concentrated, drained and used to inoculate sterilized sand (0.4-0.6mm washed aquarium sand baked at 200°C for 10 hours) in deep petri dishes in one of the three media representing the fresh, brackish and sea conditions mentioned above. Plates were placed into airtight culture containers and gassed daily as above, or left open for PAL controls. Plates were assessed for O_2_ production and redox levels after 13-15 days representing the early stationary phase in a glove box gassed with the matching culture atmosphere. Microsensors were calibrated according to the manufacturer’s instructions (UNISENSE, Aarhus, Denmark) and programmed to take readings in 50 µm increments from the surface to 6 mm beneath the surface. Each plate was measured 3-5 times depending on mat density.

### Gene screen

The genetic potential of *G. violaceus* PCC7421 (Hagemann, 2013) and *C. thermalis* PCC7203 to synthesise compatible solutes to withstand increased salt stress was determined using BLAST to identify known genes associated with salinity tolerance (Kolman & Salerno, 2015; Hagemann, 2013). In order to benefit from elevated CO_2_, cyanobacteria require a specialised transporter complement. The genomes of *G. violaceus* PCC7421 and *C. thermalis* PCC7203 were screened for the presence of CO_2_ and HCO_3_^−^ receptors (Visser *et al*., 2016) using BLAST (National Centre for Biotechnology Information, blast.ncbi.nlm.nih.gov).

### Statistics

Statistical analyses were done using the two-tailed Student´s t-test provided by Excel 2016 (Microsoft, USA) to determine the rO_2_eCO_2_ treatment influence and the effect of increasing salinities on the organisms’ growth and O_2_ production.

## Results

### Growth curves

The growth curves (Suppl. Fig. 1) indicate that both bacterial species grow better in freshwater analogous media, BG11. Interestingly, *C. thermalis* grew better when grown in ASNIII media than when grown in brackish media (BG11:ASNIII). *G. violaceus* was unable to grow in ASNIII media and this condition was therefore omitted in further experiments. Increasing salinity slows the overall increase in cell biomass. We decided to sample on days 13 – 15, late exponential/early stationary phase for both cultures, as the reproducibility was still good between culture replicates.

### Carbonate chemistry

The increased availability of Ci in the media was confirmed by the carbonate chemistry. The *p*CO2 in control media was calculated to be in the range of 833 ±35 µatm for BG11, 1113 ±84 µatm for BG11:ASNIII and 844 ±67 µatm for ASNIII medium (Table 1).

**Table 1:**
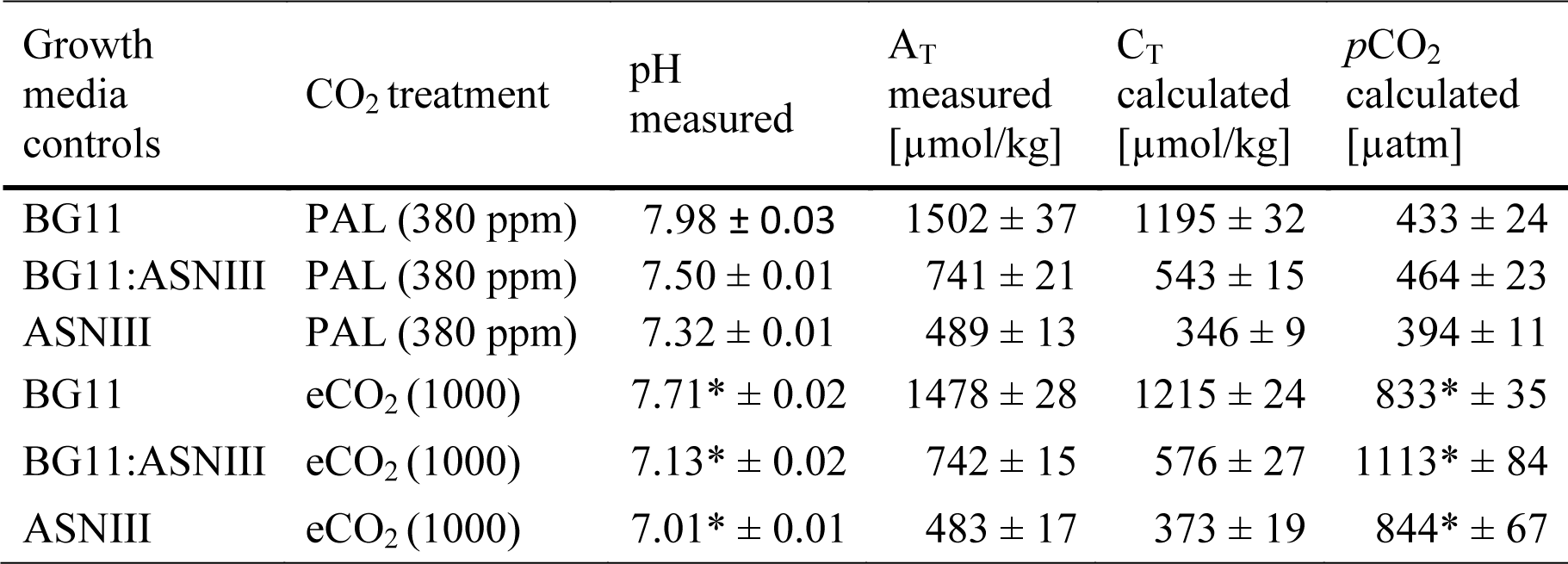
Carbonate chemistry parameter for the control media after 14 days incubation under the appropriate atmosphere either measured or calculated using CO2SYS. Statistical significant differences between the two CO_2_ treatments are indicated by * for p < 0.05 (Student´s t-test, two-tailed). Abbreviations: A_T_ total alkalinity, C_T_ total carbon.

### Salinity effects on total pigment content

Altering the salt content in media did not induce a significant change in the phycobilliprotein and chlorophyll *a* content of *C. thermalis* (Fig. 1A), however a significant reduction in carotenoid production was observed with increasing salinity. There was a significant drop in both phycobilliprotein and chlorophyll *a* content in cultures grown under rO_2_eCO_2_. Phycobilliprotein, chlorophyll *a* and carotenoid production was reduced for *G. violaceus* PCC7421 with increasing salinities and under the experimental atmosphere (Fig. 1B).

**Figure 1:**
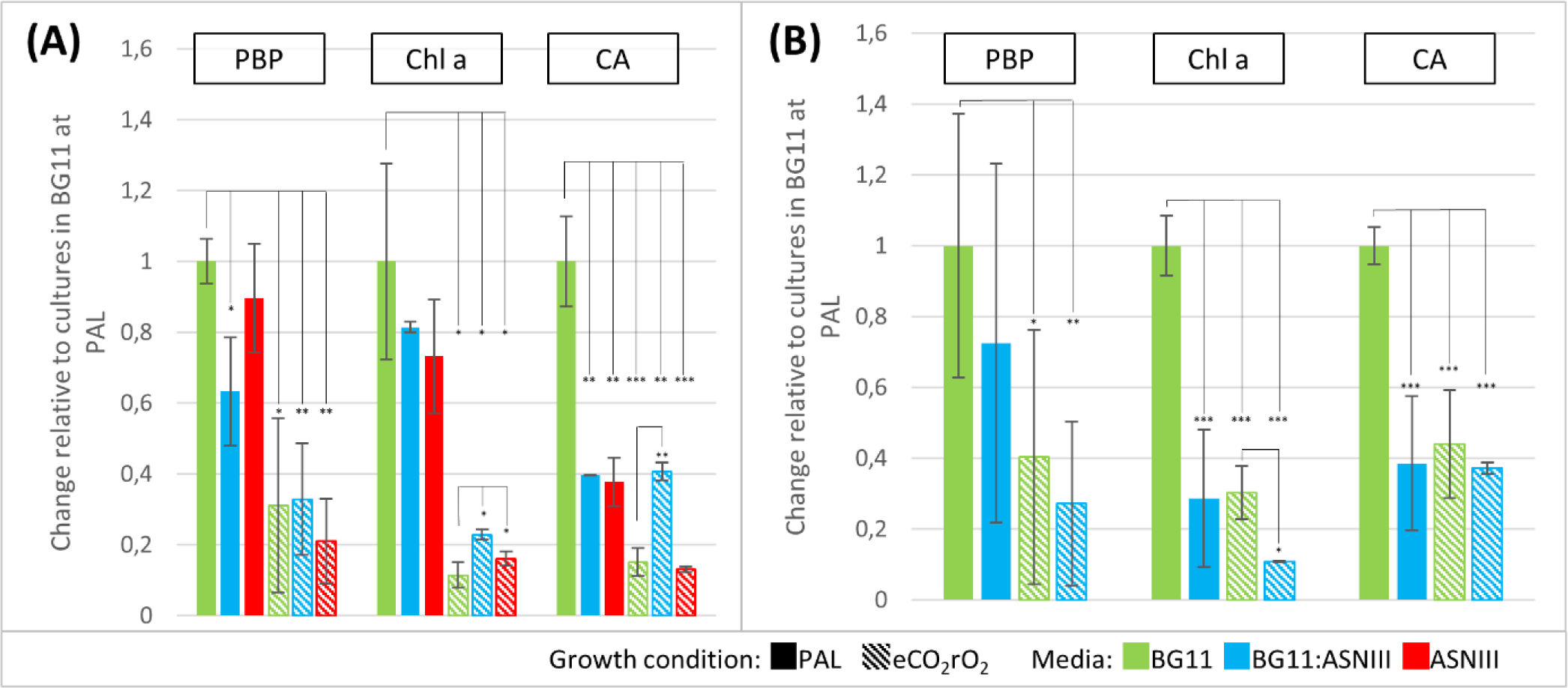
Phycobilliprotein (PBP), Chlorophyll a (Chl a) and Carotenoid (CA) content of 2 week old *C. thermalis* PCC7203 (A) and *G. violaceus* PCC7421 (B) cultures grown under different salinities and atmospheres. Pigment contend was photometrically determined in µg per ml culture from three independent samples and is expressed as relative value to respective cultures grown under PAL atmosphere in BG11 media ± SD. Cultures were incubated under PAL resp. eCO_2_rO_2_ atmosphere (solid / dashed colour) in media with increasing salinity levels from BG11 (fresh water, green) over BG11:ASNIII (brackish water, blue) to ASIII (sea water, red). Statistical significance values are indicated by * for p < 0.05; ** for p < 0.01 and *** for p < 0.001 (Student´s t-test, two-tailed)

### Salinity effects on nett photosynthesis

*Chroococidiopsis thermalis* PCC7203 showed a significant decrease in nett photosynthesis rates expressed against chlorophyll a content, with increasing salinities (Fig. 2A) when cultured under PAL or CO_2_. What is interesting is the lack of decrease in NP per µg chlorophyll when grown under a rO_2_eCO_2_ atmosphere. When expressed per ml of culture medium (Fig.2C), the NP rates are significantly reduced for all cultures grown under rO_2_eCO_2_ atmosphere compared to measurements of cultures under PAL conditions in BG11. *G. violaceus* PCC7421, grown under PAL conditions, still showed comparative NP rates per µg chlorophyll *a* to *C. thermalis* in BG11 medium (Fig. 2B). However, there was a significant decrease in NP when grown in brackish BG11:ASNIII medium or under rO_2_eCO_2_ atmosphere. Expressed as NP per ml culture (Fig.2D) cultures grown in BG:ASNIII showed a very significant decrease in CO_2_ assimilation compared to cultures in BG11 with NP close to zero if additionally incubated under a rO_2_eCO_2_ atmosphere. Additionally, no significant changes were observed in the dark respiration rates based on chlorophyll content (Suppl. Fig. 2).

**Figure 2:**
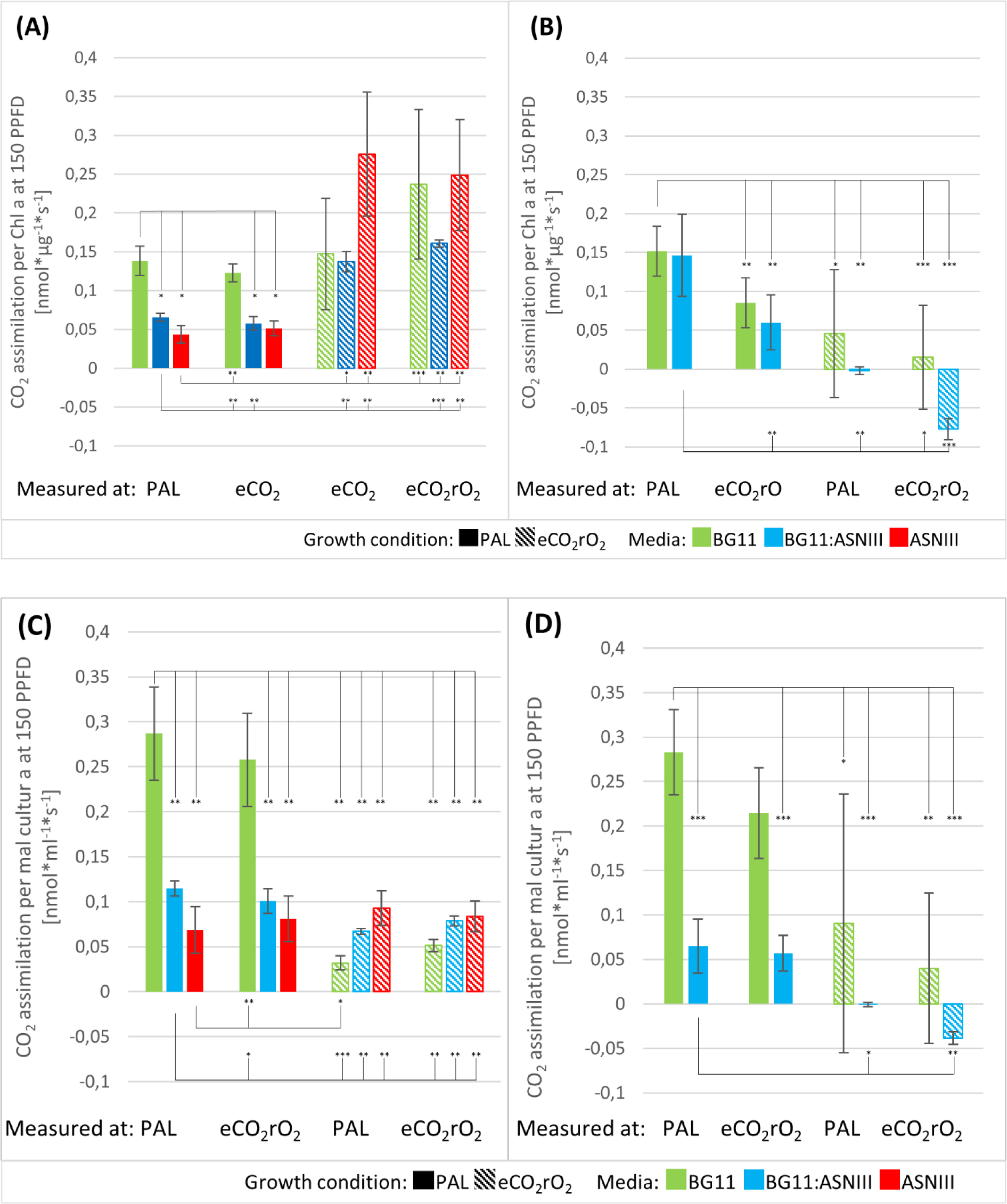
Representation of the nett photosynthesis of cultures grown under different salinities and atmospheres. The Nett photosynthesis of *C. thermalis* 7203 (A) and *G. violaceus* (B) are expressed as CO2 assimilation per second and µg Chlorophyl a, respectively per ml cultur (C) and (D). Cultures were incubated under PAL resp. eCO_2_rO_2_ atmosphere (solid / dashed colour) in media with increasing salinity levels from BG11 (fresh water, green) over BG11:ASNIII (brackish water, blue) to ASNIII (sea water, red). Statistical significance values are indicated by * for p < 0.05; ** for p < 0.01 and *** for p < 0.001 (Student´s t-test, two-tailed).

### Salinity effects on O_2_ production and redox levels in pseudomats

Microsensor readings indicated that we were indeed working under reduced O_2_ conditions, with readings approaching close to zero at 5 mm depth (Suppl. Fig. 4 B, D & E and Fig. 5 B & D). Each of the triplicate pseudomat cultures were measured a minimum of 3 times generating 9 profiles per condition (Suppl. Figs 4 & 5). The O_2_ saturation point for media varies with changing salt content, therefore, the O_2_ saturation coming from the atmosphere is indicated as grey bar of the total value displayed in Fig. 3, with coloured readings indicating actual cyanobacterial O_2_ production. The O_2_ saturation points for the rO_2_eCO_2_ atmospheres were calculated as follows: 811 nmol/L (BG11), 727 nmol/L (BG11:ASNIII) and 642 nmol/L (ASNIII). The values are too small to graphically represent in Fig. 3.

**Figure 3.**
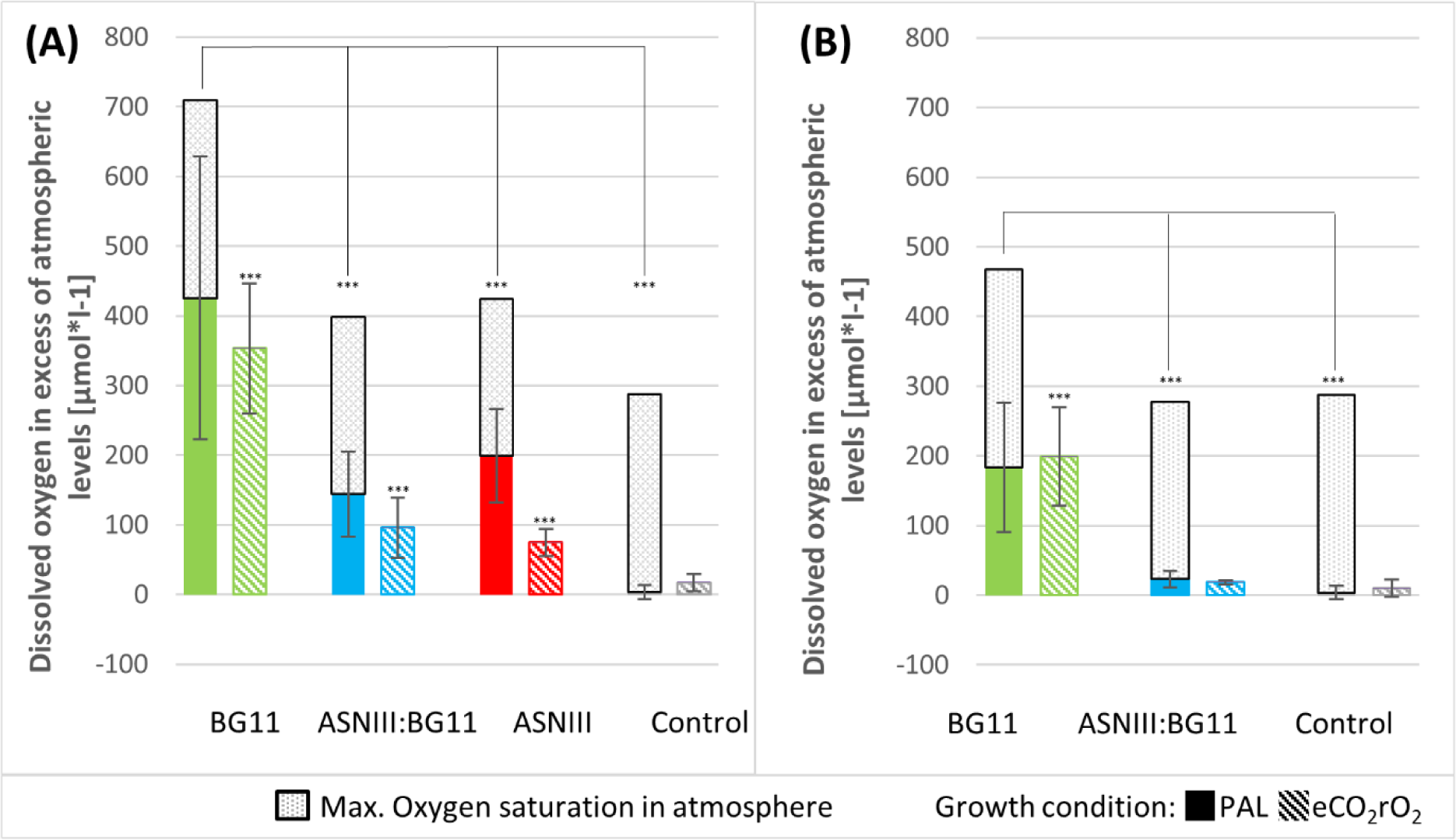
O_2_ release for A: *Chroococcidiopsis thermalis* PCC7203 and B: *Gloeobacter violaceus* PCC7421. The bars represent the average of the top 9 readings for each pseudomat profile, with their standard deviations. Each bar represents a minimum of 9 profiles (3 profiles for each triplicate for each condition). The greyed out bar indicates the O_2_ saturation derived from exposure to the atmosphere for each of the media used. The control for BG11 is also included for comparison. Statistical significance was tested for increasing salinity and different atmospheres with the same media. Values are indicated by *** for p < 0.001 (Student´s t-test, two-tailed)

*C. thermalis* PCC7203 exhibited a significant reduction in O_2_ production with increasing salinities under PAL conditions (Fig. 3), however there was still O_2_ production at higher salinity levels mimicking a marine environment. O_2_ release under a rO_2_eCO_2_ atmosphere was comparable to that under PAL for cultures growing in BG11. Increased medium salinity resulted in significant reduction of O_2_ production.

*G. violaceus* PCC7421 released comparable levels of O_2_ when grown in BG11 medium under both atmospheres tested (Fig. 3), but was significantly lower than recorded for *C. thermalis* PCC7203. Growing the pseudomats in brackish water significantly reduced the O_2_ release to the environment, however some O_2_ production was evident above the background within the first 5 mm under PAL conditions, with no O_2_ release observed under a rO_2_eCO_2_ atmosphere (Suppl. Fig. 4).

The overall reduction in redox potential of the environment with reduced O_2_. No real changes of redox values were observed in profiles through the pseudomats. The average of the top ten values are plotted below (Fig. 4) with those of the uninoculated control plates for comparison. The redox values recorded for *C. thermalis* PCC7203 grown under PAL conditions show a comparable increase for all three media tested, when compared to the redox readings obtained for the control uninoculated plates. No significant change in redox otential was observed for pseudomats grown under a rO_2_eCO_2_ atmosphere although the average reading was reduced in BG11 and BG11:ASNIII medium, suggesting an altered redox balance (Fig. 4). The redox readings for *G. violaceus* pseudomats grown under freshwater conditions were raised compared to control values, and equalled those observed for *C. thermalis* cultured in BG11 (Fig. 4). When grown under rO_2_eCO_2_, *G. violaceus* pseudomats showed lower average redox values than were recorded for the control media.

**Figure 4:**
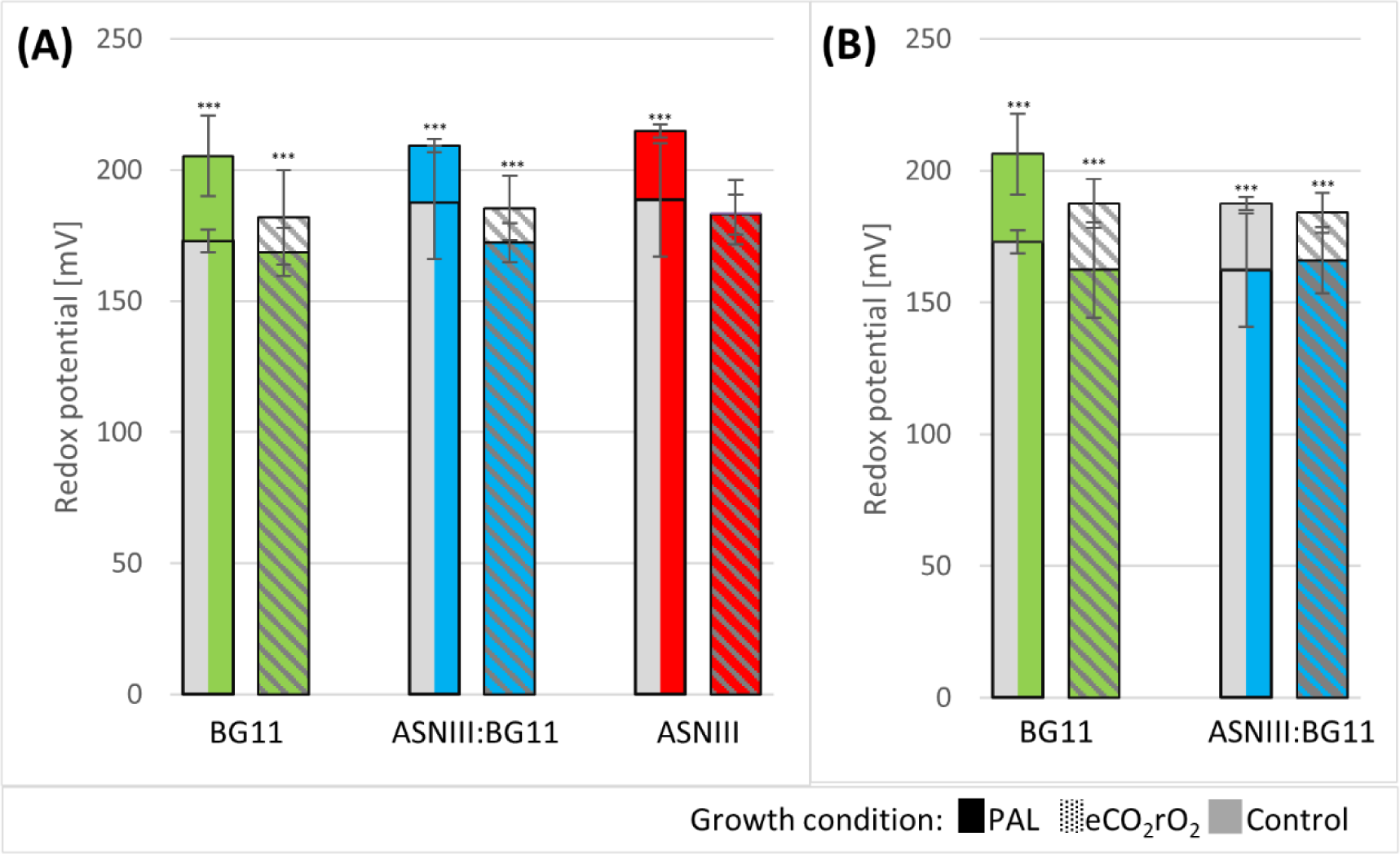
Redox potential profile for pseudomats of A: *Chroococcidiopsis thermalis* PCC7203 and B: *Gloeobacter violaceus* PCC7421. The coloured bars represent the average of the top 10 readings within the first 200 µm for each pseudomat profile, with their standard deviations. Each bar represents a minimum of 9 profiles (3 profiles for each triplicate for each condition). The control values for each media are indicated as grey bars / grey dashed area. Statistical significance was tested between inoculated media and control. Values are indicated by *** for p < 0.001 (Student´s t-test, two-tailed).

**Figure 5:**
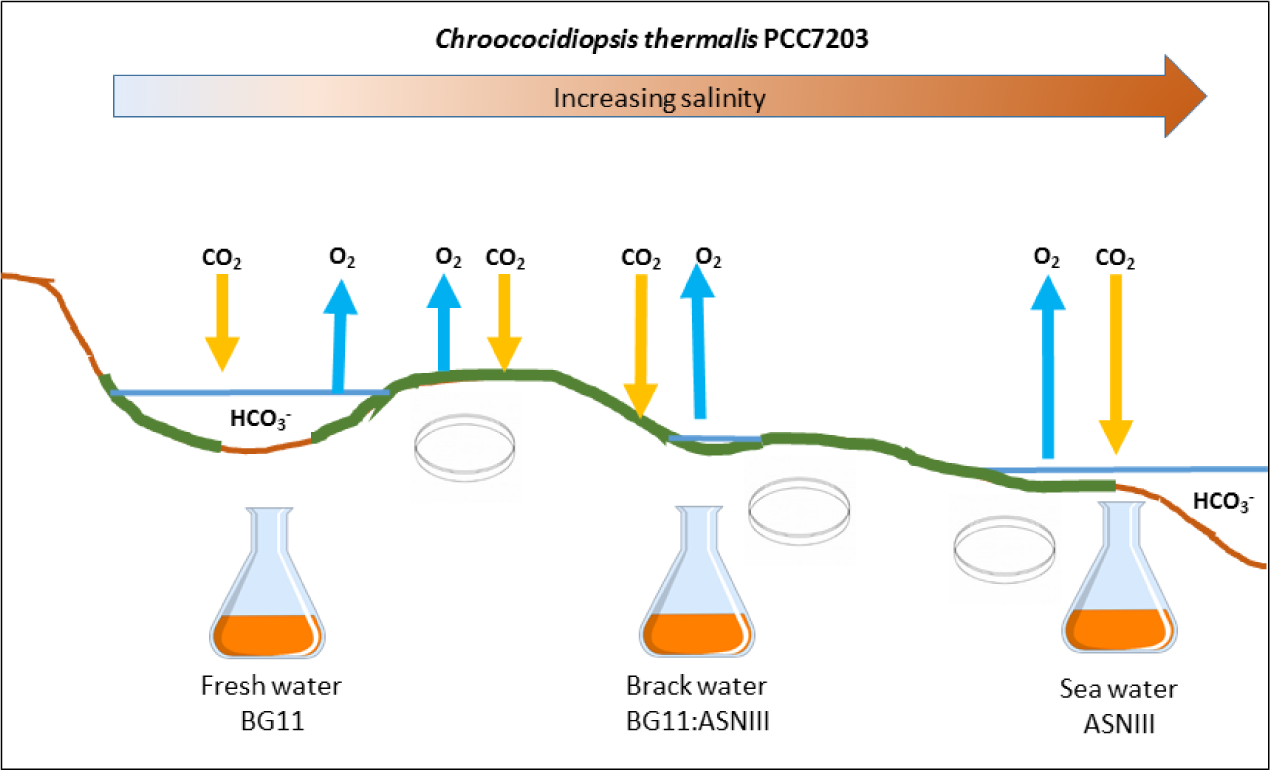
Graphical representation of the transfer of an ancestral freshwater cyanobacterial species through increasing salinities during the Archaean. Please see text for a complete explanation. CO_2_ with yellow arrow represents photosynthetic activity (measured by gas exchange in this study). O2 with a blue arrow represents the release of oxygen resulting from active photosynthesis (measured using O_2_ microsensors in this study). The flasks represent the liquid cultures and the plates represent the pseudomats used to assess cyanobacterial growth under increasing salinities from fresh, through brackish to marine strength salinity

### Genetic acclimation potentential

*Gloeobacter violaceus* PCC 7421 was previously shown to carry the genes for sucrose-phosphate synthase (Hageman 2013). Our screen (Table 2) confirmed this and identified two additional Na^+^/H^+^ antiporter receptors related to salt tolerance (Inaba et al., 2001) encoded on the genome of *G. violaceus* PCC7421. A screen of the *C. thermalis* PCC7203 genome revealed a similar salt tolerance gene profile with the addition of a high flux Na^+^/H^+^ antiporter receptor and the genes for trehalose synthesis. The C_i_ uptake receptor profile for both organisms indicates that they carry the genes for both gaseous CO_2_ uptake receptors and the 2 high specificity/low flux HCO3-receptors. Significantly, they also carry the gene encoding the low specificity/high flux receptor, BicA, conferring on both strains the ability to benefit from increased levels of HCO_3_ ^−^ in the media resulting from elevated atmospheric CO_2_ levels.

**Table 2:** Gene screen results for *C. thermalis* PCC7203 and *G. violaceus* PCC7421. The genomic sequences of both organisms were screened for the presence of genes conferring salt tolerance and the receptors involved in C_i_ uptake. The presence or absence of the gene is indicated by a + or – for each genome and the gene description follows in the final column. Results marked with an ¹ are from (Kolman et al., 2015) and ² from Hageman et al., (2013). Details on tested sequences and E-values are described in Supply. Tab.1.

| Protein | *C. thermalis*<br>PCC7203 | *G. violaceus*<br>PCC7421 | Description |
| --- | --- | --- | --- |
| SpsA | +<sup>1</sup> | +<sup>2</sup> | Sucrose-phosphate synthase |
| Spp | +<sup>1</sup> | +<sup>2</sup> | Sucrose-phosphate phosphatase |
| OtsAB | - | -<sup>2</sup> | Trehalose-6-phosphate synthase/HAD<br>hydrolase |
| TreZ | + | -<sup>2</sup> | Malto-oligosyltrehalose<br>trehalohydrolase |
| TreY | + | -<sup>2</sup> | Malto-oligosyltrehalose synthase |
| TreA | - | -<sup>2</sup> | Alpha,alpha-trehalase |
| GgpS | - | -<sup>2</sup> | Glucosylglycerol-phosphate synthase |
| GgpP | - | -<sup>2</sup> | Glucosylglycerol 3-phosphatase |
| GpgS | - | -<sup>2</sup> | Glucosylglycerol phosphate synthase |
| GgpgP | - | -<sup>2</sup> | Glucosyl-3-phosphoglycerate<br>phosphatase |
| GsmT | - | -<sup>2</sup> | Glycine/sarcosine N-methyltransferase |
| DmT | - | -<sup>2</sup> | Sarcosine/dimethylglycine N-methyltransferase |
| NhaS1 | + | - | Low-affinity Na(+)/H(+) antiporter |
| NhaS3 | + | + | High-flux Na+/H+ antiporter |
| NhaS4 | + | + | Na+/H+ antiporter |

| Protein | *Chroococcidiopsis thermalis* PCC 7203 | *Gloeobacter violaceus* PCC 7421 | Description |
| --- | --- | --- | --- |
| BicA | + | + | Low-affinity bicarbonate/sodium symporter |
| SbtA | + | - | High-affinity bicarbonate/sodium symporter |
| CmpA | + | + | High-affinity ATP-dependent bicarbonate uptake system |
| CmpB | + | + | High-affinity ATP-dependent bicarbonate uptake system |
| CmpC | + | + | High-affinity ATP-dependent bicarbonate uptake system |
| CmpD | + | + | High-affinity ATP-dependent bicarbonate uptake system |
| NadF3 | + | + | High-affinity CO<sub>2</sub> uptake system |
| NadhD3 | + | + | High-affinity CO<sub>2</sub> uptake system |
| NdhD4 | + | + | Low-affinity CO<sub>2</sub> uptake system |
| NdhF4 | + | + | Low-affinity CO<sub>2</sub> uptake system |
| ChpY | + | + | High-affinity CO<sub>2</sub> uptake system |
| ChpX | + | + | Low-affinity CO<sub>2</sub> uptake system |
| NahaS3 | + | + | Sodium/proton antiporter |
| CcmR2 | + | + | Regulator of sodium-dependent bicarbonate uptake operon |
| CcmR | + | + | Regulator of the high-affinity CO<sub>2</sub> uptake operon |

## Discussion

Recently publications have provided evidence that oxidative photosynthesis, namely the splitting of water, evolved prior to the establishment of the oxidative phototrophs, cyanobacteria (Cardona, 2015, Cardona *et al*., 2015). This, combined with evolutionary studies indicating that the crown cyanobacteria evolved prior to the great oxygenation event in freshwater environments (Sanchez-Baracaldo, 2015, Sanchez-Baracaldo *et al*., 2017) has altered our understanding of how the world as we know it evolved: specifically regarding the conditions leading up to the GOE. First and foremost is the question of whether cyanobacteria could not just survive, but grow under increasing salinities to eventually give rise to the free ocean dwelling species thought to have precipitated the GOE 2.33 Ga. This study aimed to investigate the photosynthetic ability of the root cyanobacterial species, *G. violaceus* PCC7421 and a known salt tolerant ancient lineage cyanobacterium, *C. thermalis* PCC7203 to increasing salinities under present atmospheric conditions and those thought to have prevailed towards the end of the Archaean era.

*C. thermalis* PCC7203, while showing a significant decrease in overall NP activity with increasing salinities, was still able to grow and sequester CO_2_ when cultured in a PAL atmosphere (Fig. 2A & C). *G. violaceus* PCC7421 significantly reduced its NP levels when grown in brackish medium under PAL atmospheric conditions (Fig 2B) and died in the sea salt analogous medium, ASNIII. While both organisms carried the genes for sucrose phosphate synthase, *C. thermalis* can potentially express genes for trehalose synthesis as well as an additional low affinity Na^+^/H^+^ antiporter receptor (Table 2), thereby providing it with an additional potential mechanism to overcome salt stress.

Both organisms showed reduced levels of NP when grown under the rO_2_eCO_2_ atmosphere (Fig. 2 A & B). Interestingly, the NP expressed per µg chlorophyll *a* or per ml of culture showed no reduction for *C. thermalis* with increasing salinity when compared to the fresh water control (Fig. 2C). The overall reduction in NP observed in cultures grown under the rO_2_eCO_2_ atmosphere is presumably the result of reduced pigment levels (Fig 1 A & B). The presence of the low specificity / high flux HCO_3_^−^ receptor gene, *bic*A, would suggest that both strains should benefit from elevated HCO_3_^−^ levels in the media resulting from eCO_2_ in the atmosphere (Visser *et al*., 2016). This was not observed in our study and may rest on several factors: A) the receptor is not expressed, B) the reduced O_2_ levels restricted oxygen dependent pathways within the organisms, thereby restricting growth or C) the CO_2_ was rapidly utilised in the closed system and the cells were Ci starved. While the control media incubated at the rO_2_eCO_2_ atmosphere contained elevated levels of C_i_ (Table 1), the C_i_ of the cell culture media was not determined. This is currently under investigation.

While significant differences were observed for NP rates determined per ml of culture under the two different atmospheres, the O_2_ production rates appeared to be similar for each salinity tested (Fig 3), regardless off the atmosphere under which it was cultured. redox levels were increased in all cultures grown under PAL when compared to the control media, except for *G. violaceus* grown under brackish conditions (Fig. 4). In contrast, all cultures grown under the rO_2_eCO_2_ atmosphere showed redox values lower than the control media. This apparent reduction was however not significant.

While the exact nature of the rO_2_eCO_2_ increase in Ci availability in the cultures remains to be determined, we are still able to make some clear observations from this study. We have demonstrated that the fresh water strain of *C. thermalis* PCC7203 continues to fix CO_2_ at salinities representing brackish and sea salinities under PAL conditions. While its survival was previously demonstrated up to marine salinity of 250 mM NaCl (Cumbers & Rothschild, 2014), the physiological effects of elevated salt on the process of oxidative photosynthesis had not been determined. Transfer of *C. thermalis* into brackish and sea salt media resulted in significant reduction in NP levels, presumably as the result of reduced pigment synthesis resulting from salt stress. *G. violaceus* in contrast was barely able to continue with NP in a brackish environment and not at all able to grow in marine analogous media (not shown). The additional genetic ability to synthesise trehalose presumably conferred upon *C. thermalis* an advantage to *G. violaceaus*, allowing it to grow in sea water.

Repeating this experiment under a rO_2_eCO_2_ atmosphere also showed that the two strains could continue photosynthesising in freshwater conditions, albeit at reduced levels compared to present levels of CO_2_ and O_2_ in the atmosphere. *C. thermalis* could also continue photosynthesising under increased salinities with comparable NP activities per µg chlorophyll *a* in this oxygen poor environment (Fig. 2). Overall biomass was reduced in the rO_2_eCO_2_ cultures as evidenced by the reduced pigment content per ml of culture (Fig. 1). The lack of increased respiration rates (Suppl. Fig. 2) and carotenoid synthesis (Fig. 1) suggests that even under increased salinities, the *C. thermalis* PCC7203 cultures were not overly stressed.

Taken together, this data supports the hypothesis of Cyanobacterial evolution in freshwater environments and their transition into increasingly salty environments (Sanchez-Baracaldo, 2015, Sanchez-Baracaldo *et al*., 2017) as illustrated in Fig. 5. Cyanobacteria originated in a freshwater environment which would also have allowed the formation of loose microbial mats on solid substrates along the edges of the water body. Occasional flooding events would have transferred cyanobacterial biomass along a downward gradient, allowing them to colonise potentially brackish environments, possibly in a delta scenario. Further flooding from either an ocean or wash out from inland freshwaters, could have introduced the originally freshwater strain into an ocean environment. By demonstrating active photosynthesis in both liquid cultures and pseudomats, we have shown that some cyanobacterial species, equipped with the minimal gene complement of sucrose and trehalose synthesis, would have been able to make the gradual transition into increasingly saline environments, not only in PAL atmospheric conditions, but under the reduced oxygen and elevated CO_2_conditions thought to have existed during the late Archaean.

## Supplementary data

**Supplementary Figure 1:**
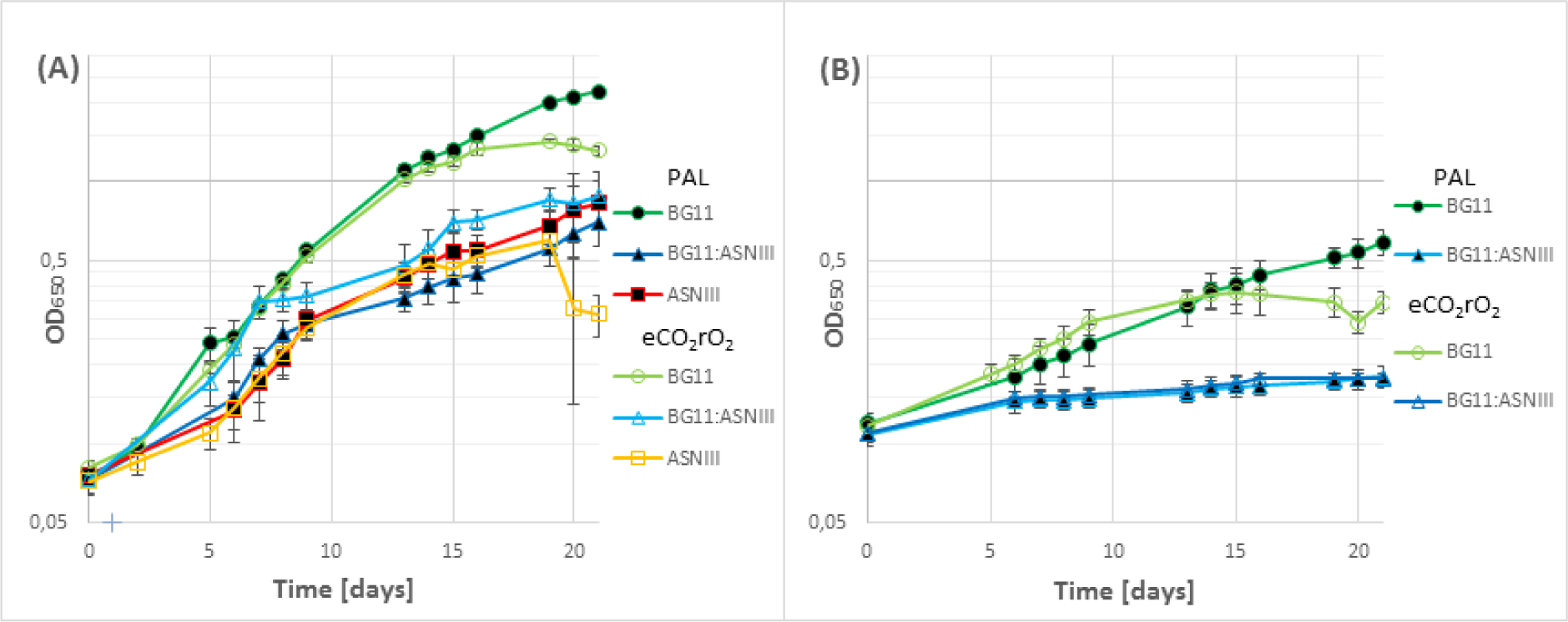
Cyanobacterial growth curves under increasing salinities for **A:** *Chroococcidiopsis thermalis* PCC7203 and **B:** *Gloeobacter violaceus* PCC7421. All cultures were exposed to a PAL atmosphere for 5-10 minutes at every absorbance reading.

**Supplementary Figure 2:**
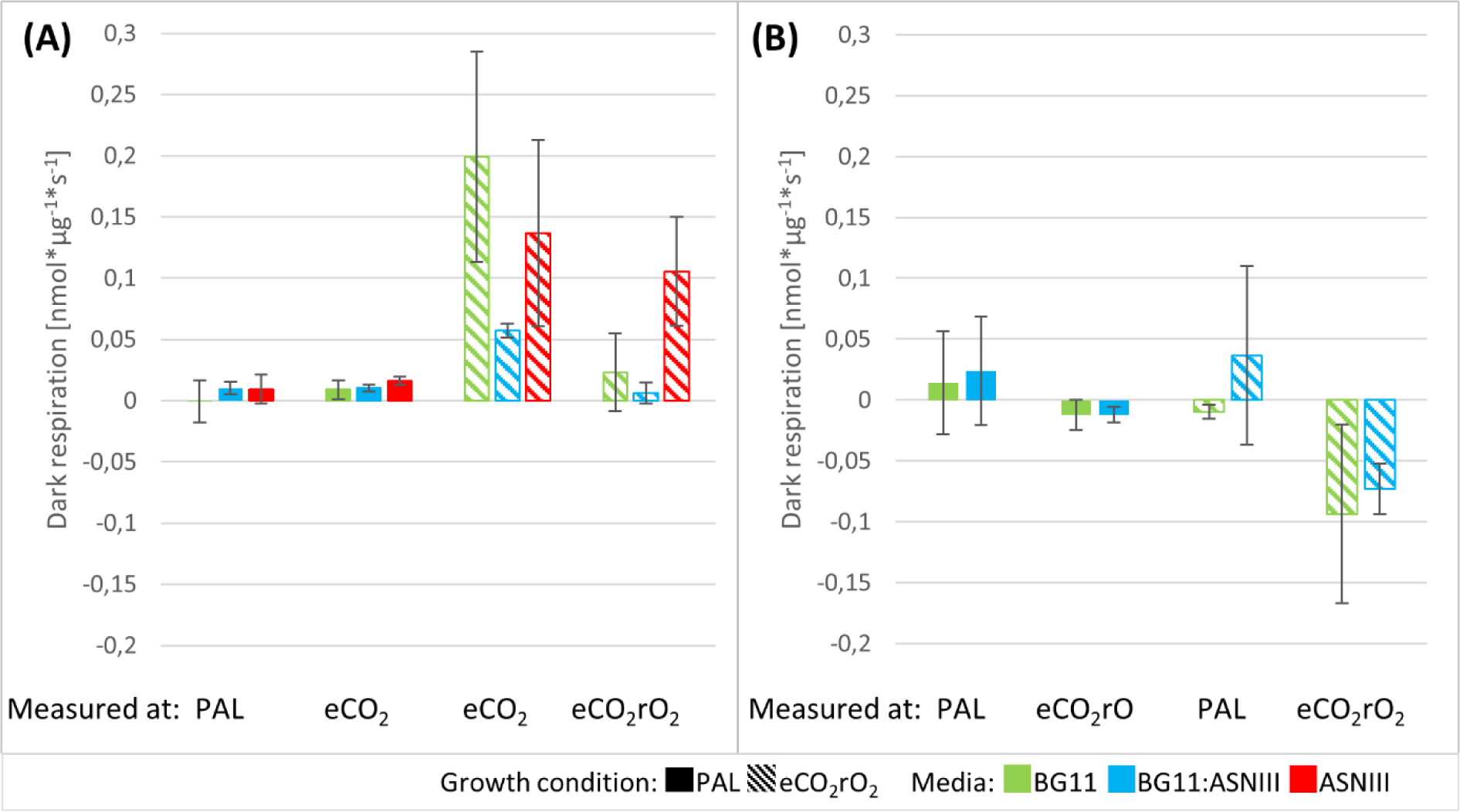
Representation of the dark respiration rates of cultures grown under different salinities and atmospheres. Respiration rates expressed for chlorophyll *a* content of *C. thermalis* 7203 (A) and *G. violaceus* (B) with increasing salinities and rO_2_eCO_2_ conditions. *G. violaceus* cultures grown under PAL conditions and in freshwater were also assessed for their NP under eCO_2_rO_2_.

**Supplementary Figure 3:**
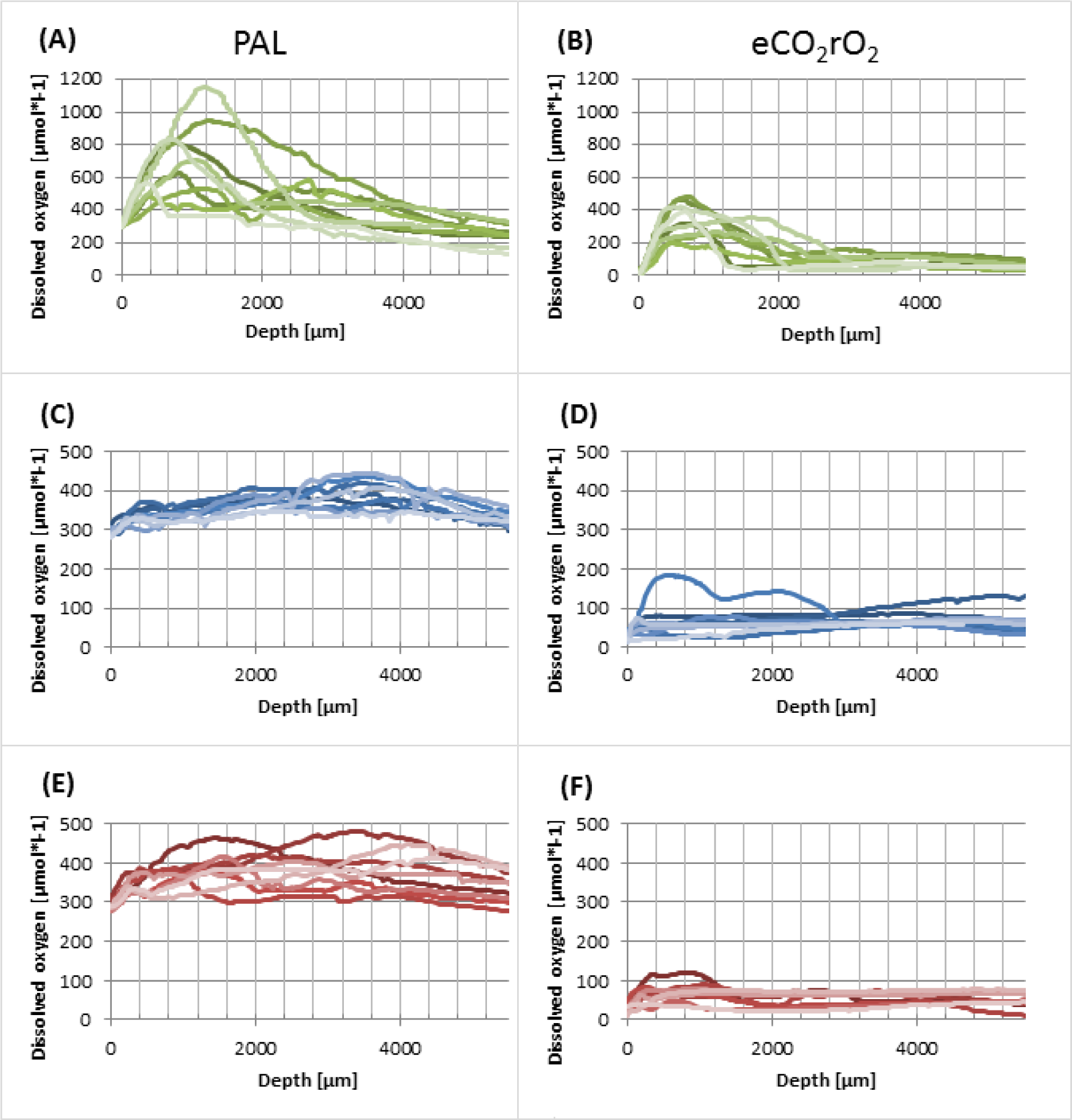
Dissolved oxygen measured in pseudomats of *Chroococcidiopsis thermalis* PCC7203. A, C, E, are grown under PAL conditions while B, D and F are grown under rO_2_eCO_2_. Pseudomats A & B are grown in freshwater analogous medium BG11, C & D in brackish analogous medium BG11:ASNIII while E & F represent pseudomats grown in sea water analogous medium ASNIII. First surface contact was determined as readings over or under atmospheric levels.

**Supplementary Figure 4:**
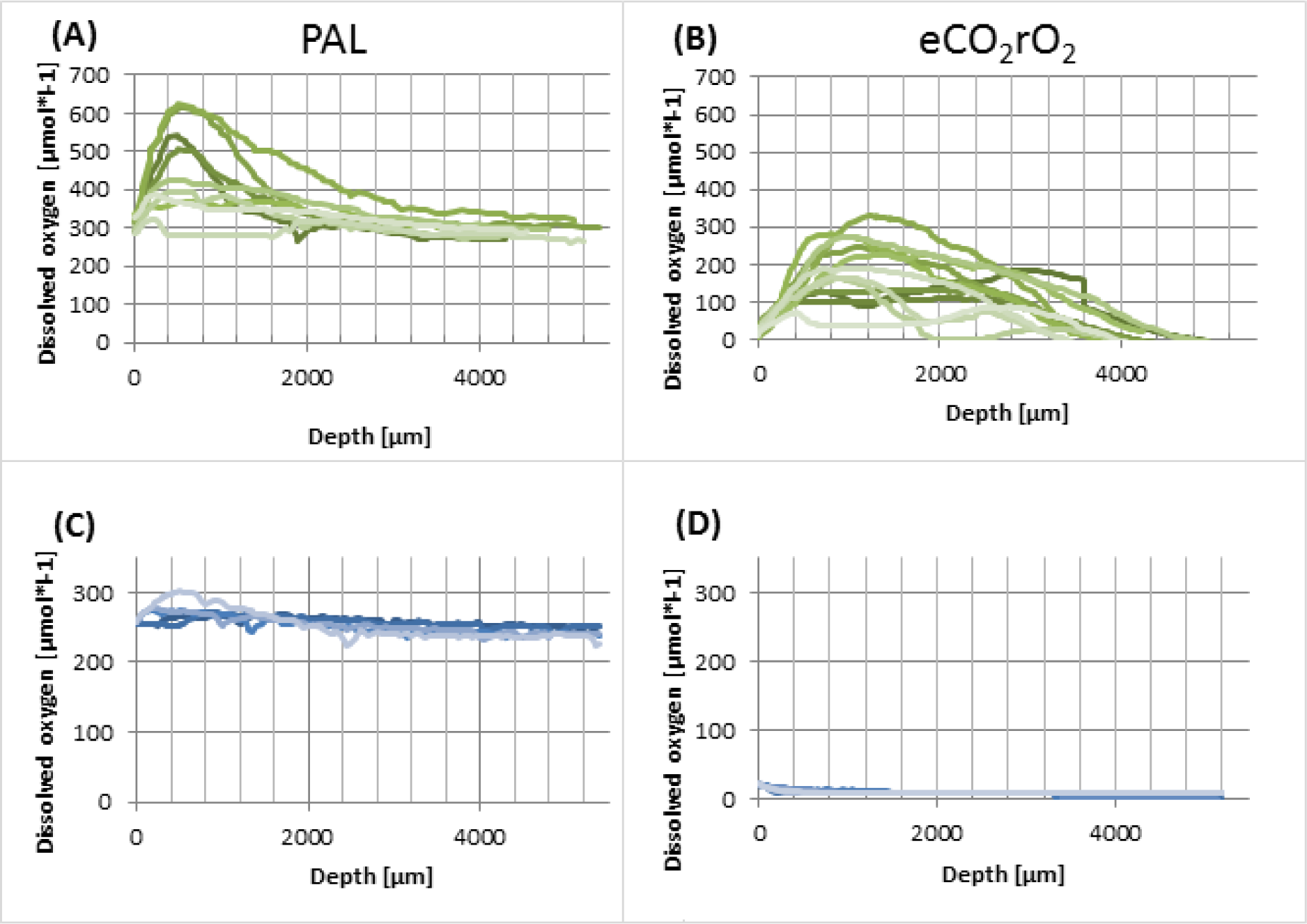
Dissolved oxygen measured in pseudomats of *Gloeobacter violaceus* PCC7421. A &C are grown under PAL conditions while B & D are grown under rO_2_eCO_2_. Pseudomats A & B are grown in freshwater analogous medium BG11, while C & D represent pseudomats grown in brackish analogous medium BG11:ASNIII.

**Supplementary Table 1:**
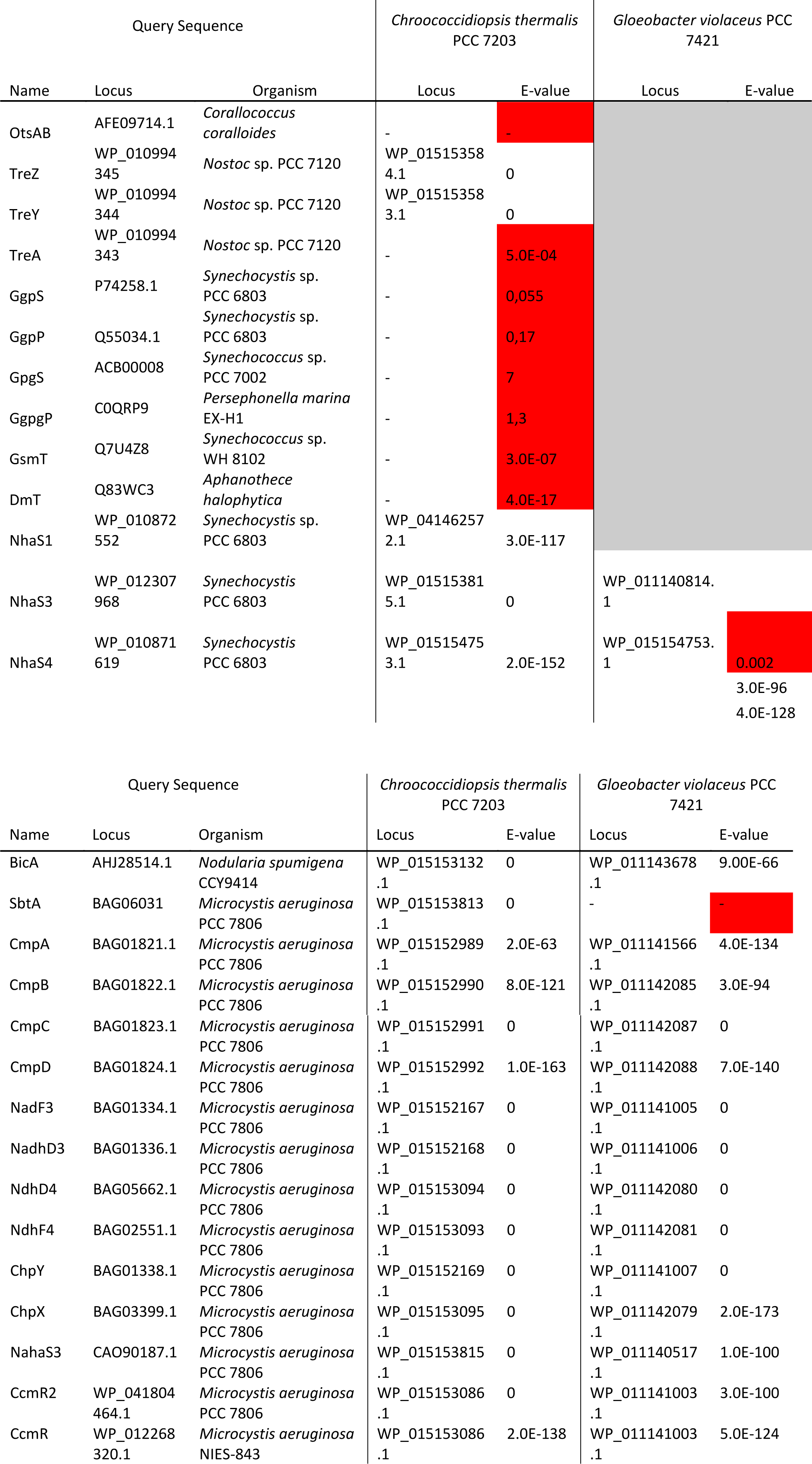
Gene screen results for *C. thermalis* PCC7203 and *G. violaceus* PCC7421. The genomic sequences of both organisms were screened for the presence of genes conferring salt tolerance and the receptors involved in C_i_ uptake. The gene ID and name used for the screen is listed under the Protein column. High E-values, indicating a random hit, or no matches at all are marked in red.

